# Post-hoc long-read sequencing links leukemic mutation status to single-cell transcriptomes

**DOI:** 10.64898/2026.05.06.723232

**Authors:** Sofia Papavasileiou, Chenyan Wu, Daryl Boey, Lucille Margerie, Jiezhen Mo, Ulla Olsson-Strömberg, Stina Söderlund, Gunnar Nilsson, Joakim S. Dahlin

## Abstract

Single-cell RNA-sequencing-based characterization of cells that belong to the neoplastic clone is a major challenge in hematologic neoplasms, where malignant and normal cells coexist. Confident molecular profiling requires simultaneous analysis of gene expression and genetic mutations in individual cells, an ability that is not supported by the standard 10X Genomics workflow. Here, we systematically evaluated the potential and limitations of repurposing amplified cDNA generated during the 10X Genomics 3’ workflow for post-hoc genotyping of individual cells. We first established a mixed leukemic cell line system comprising one cell line with *KIT* point mutations and another with the *BCR::ABL1* fusion gene. Targeted long-read PacBio sequencing enabled post-hoc assignment of mutation data to transcriptionally profiled cells, but recovery differed between targets. Consistent with ambient RNA in microfluidics-based single-cell workflows, mutation-associated transcripts were detected in cells not expected to carry the corresponding mutations, illustrating how transcript recovery complicates cell-level genotype assignment. Target-specific thresholds mitigated this source of misclassification. In primary chronic myeloid leukemia samples, the post-hoc approach detected *BCR::ABL1*-positive cells at diagnosis, but not during imatinib treatment. Together, we present a framework for adding mutation status to cells already profiled using the 10X Genomics workflow and highlight broader considerations for transcript-based single-cell genotyping.

**Novelty Statements:** *What is the NEW aspect of your work?:* We present and evaluate a novel framework for post-hoc detection of leukemic mutations in cells profiled with the 10X Genomics 3’ single-cell RNA-sequencing workflow, thereby adding mutation status to single-cell transcriptomes.

*What is the CENTRAL finding of your work?:* The framework identifies mutation-carrying cells within heterogeneous single-cell RNA- sequencing datasets, addressing a practical challenge in the analysis of patient-derived samples where relevant mutations were not known or not targeted as part of the original transcriptomics profiling.

*What is (or could be) the SPECIFIC clinical relevance of your work?:* Mutation-informed characterization of cells from patients with hematologic neoplasms has the potential to support the identification of clinically relevant cell populations associated with treatment response or disease progression.

## Introduction

Somatic mutations are associated with many hematologic neoplasms. For example, the somatic *BCR::ABL1* gene fusion is the cause of chronic myeloid leukemia [1], and a point mutation in the *KIT* gene drives the clonal mast cell accumulation in systemic mastocytosis and mast cell leukemia [2, 3]. High-throughput single-cell RNA-sequencing technologies now allow for large-scale profiling of the patients’ hematopoietic cells, which comprise a mixture of mutant and wild-type cells. Thus, specific analysis of the mutated cells’ transcriptome requires simultaneous genotyping to be performed. In theory, one can detect somatic mutations at the RNA level. In practice, high-throughput single-cell RNA- sequencing platforms such as the one from 10X Genomics are designed to capture the 5’ or 3’ ends of individual transcripts using Illumina sequencing. This inherently complicates the detection of mutations distant from the transcript ends.

Genotyping of Transcriptome (GoT) is an elegant approach that detects mutations far from the 3’ end of transcripts by customizing the 10X Genomics-based experimental pipeline [4]. The GoT approach includes preselection of genes of interest and preferential amplification of their transcripts as part of the 10X workflow. This is followed by specialized molecular processing that enables mutation detection by short-read sequencing. However, the relevant targets may not be known when diagnostic patient samples are processed using the 10X workflow. This motivates the development and evaluation of post-hoc mutation detection approaches that repurpose the stable amplified cDNA pool generated without gene-specific enrichment.

Long-read sequencing is an approach to determine sequence variations distant from the transcript ends [5, 6]. However, performing long-read sequencing on the amplified cDNA pool inherently biases the analysis to highly expressed genes. Single-cell Targeted Isoform Long-Read sequencing (scTaILoR-seq) and long-read genotyping of transcripts (LOTR-seq) use panels to enrich for specific genes before long-read sequencing [7–9], facilitating the detection of single nucleotide variants among predefined subsets of genes. To genotype leukemic cells, it is often relevant to specifically target patient-specific mutations. Consequently, a method for post-hoc, targeted genotyping of leukemic mutations was recently developed for 5’-based scRNA-seq, utilizing Nanopore-based long-read sequencing [10].

Here, we systematically evaluated the potential and limitations of post-hoc genotyping of individual cells processed using the 10X Genomics 3’ scRNA-seq platform. The method is established through a carefully controlled setup, in which two leukemic cell lines with distinct genotypes are mixed and subjected to single-cell RNA-sequencing. By repurposing a stable intermediate product of the 10X Genomics workflow, we show that leukemic mutations are detected using long-read sequencing. Specifically, *BCR::ABL1* fusion transcripts and point mutations in *KIT* are captured in individual cells. We apply the method on diagnostic and treatment samples of chronic myeloid leukemia, showing the loss of *BCR::ABL1* signal upon treatment. Taken together, we provide a method for post-hoc genotyping of cells processed using the 10X Genomics 3’ platform and highlight broader considerations for transcript-based genotyping of single cells.

## Methods

### Single-cell RNA-sequencing data generation

HMC1.2 and Ku812 cell lines were hashtagged with oligonucleotide-labeled antibodies (TotalSeq-B0253 or TotalSeq-B0254; Biolegend, San Diego California, USA) before mixing and performing single-cell RNA-sequencing. Leukocytes from a patient with chronic myeloid leukemia were isolated at diagnosis and during imatinib treatment using BD Pharm Lyse (BD Biosciences, Franklin Lakes, New Jersey, USA). For details related to the culture of the cell lines and leukocyte isolation, see the Supplementary Methods.

The cells were processed using the Chromium Next GEM Single Cell 3’ Reagent Kit v3.1 (dual index) (10X Genomics, Pleasanton, California, USA). For the cell line-related datasets, the single-cell transcriptomics workflow was combined with the Feature Barcode technology for Cell Surface Protein to capture the hashtag oligonucleotide sequences and thereby distinguish the two cell lines. For all datasets, the standard scRNA-seq workflow was adapted to favor the capture of longer cDNA products. Specifically, the reverse transcription step was performed for 2 hours instead of 45 minutes and the extension time during the first cDNA amplification was increased from 1 minute to 3 minutes [5]. These extended reaction times are expected to increase the recovery of longer transcript fragments. The short-read sequencing libraries were sequenced on the Illumina platform.

### Targeted amplification of BCR::ABL1, BCR and KIT cDNA

Q5 HotStart High-Fidelity MasterMix (New England Biolabs, Ipswich, Massachusetts, USA) was used in all the PCR reactions performed to amplify the genes of interest and the PCR products were size selected, and gel purified using the Wizard SV Gel and PCR Clean-Up system (Promega, Madison, Wisconsin, USA). The final primer concentration in all the reactions was 1 μM. All the primers used in this study were purchased from Sigma-Aldrich (St. Louis, Missouri, USA). The annealing temperatures and extension times were adjusted according to the primers used and the length of the target molecule (Supplementary Tables E1 and E2).

Briefly, 5 μl of the PCR-amplified cDNA generated during the 10X scRNA-seq workflow was used as input for the first amplification reactions using the universal reverse primer (containing the Read 1 and an overhang sequence [11]), and one of the mutation specific forward primers (Supplementary Table E2). Three independent PCR reactions with unique forward primers were performed to amplify 1) wild-type *BCR* and *BCR::ABL1*, 2) *KIT* c.1679T>G, and 3) *KIT* c.2447A>T. Amplified cDNA from each reaction was purified using agarose gel electrophoresis followed by excision of fragments corresponding to the amplification products. The recovered cDNA from each reaction was used as input for a second round of semi-nested PCR amplifications using the reverse universal primer against the overhang and one of the corresponding fragment-specific nested forward primers (Supplementary Table E2). The PCR products were then purified using SPRI-select beads (Beckman Coulter, Brea, California, USA) and were subjected to one additional amplification and purification round to ensure enough obtained material for the generation of long-read sequencing libraries.

Prior to long-read sequencing, gel electrophoresis confirmed the length of the amplified molecules, and the fragment identity was verified using Sanger sequencing. For the cell line-related dataset, the molecules obtained from each of the three amplification reactions were pooled together and used as input for the generation of one long-read sequencing library (Pacific Biosciences, California, USA). Targeted amplification of wild-type *BCR*/*BCR::ABL1* was performed for the patient samples. Each sample (collected before or during imatinib treatment) was used for the generation of an independent long-read sequencing library. The long-read libraries were sequenced on the Revio instrument. For details related to the library preparation and sequencing, see Supplementary Methods.

### Bioinformatics analysis

Data processing is detailed in Supplementary Methods. Quality control filtering thresholds were applied as described in Supplementary Table E3. PacBio sequencing and read processing statistics are summarized in Supplementary Tables E4-E5. Long-read sequencing-based genotyping was performed using wild-type and mutant reference sequences detailed in Supplementary Table E6, with classification thresholds described in Supplementary Table E7.

### Use of ChatGPT

ChatGPT (OpenAI) was used to improve the readability of the manuscript. The authors carefully reviewed and edited the AI-assisted text and take full responsibility for the contents.

## Results

### The 10X Genomics scRNA-seq workflow poorly detects common leukemic mutations

To evaluate whether the 10X Genomics workflow allows detection of common leukemic mutations, we performed single-cell RNA-sequencing of two unrelated cell lines: 1) the Ku812 cell line [12], which harbors the *BCR::ABL1* gene fusion that drives chronic myeloid leukemia, and 2) the HMC1.2 cell line that exhibits two point mutations in the *KIT* gene that drive systemic mastocytosis and mast cell leukemia [13]. We hashtagged the cell lines with unique oligonucleotide-conjugated antibodies before cell line mixing (Figure 1A), allowing confident annotation of each single cell’s identity in the resulting dataset (Supplementary Figure E1A).

**Figure 1.**
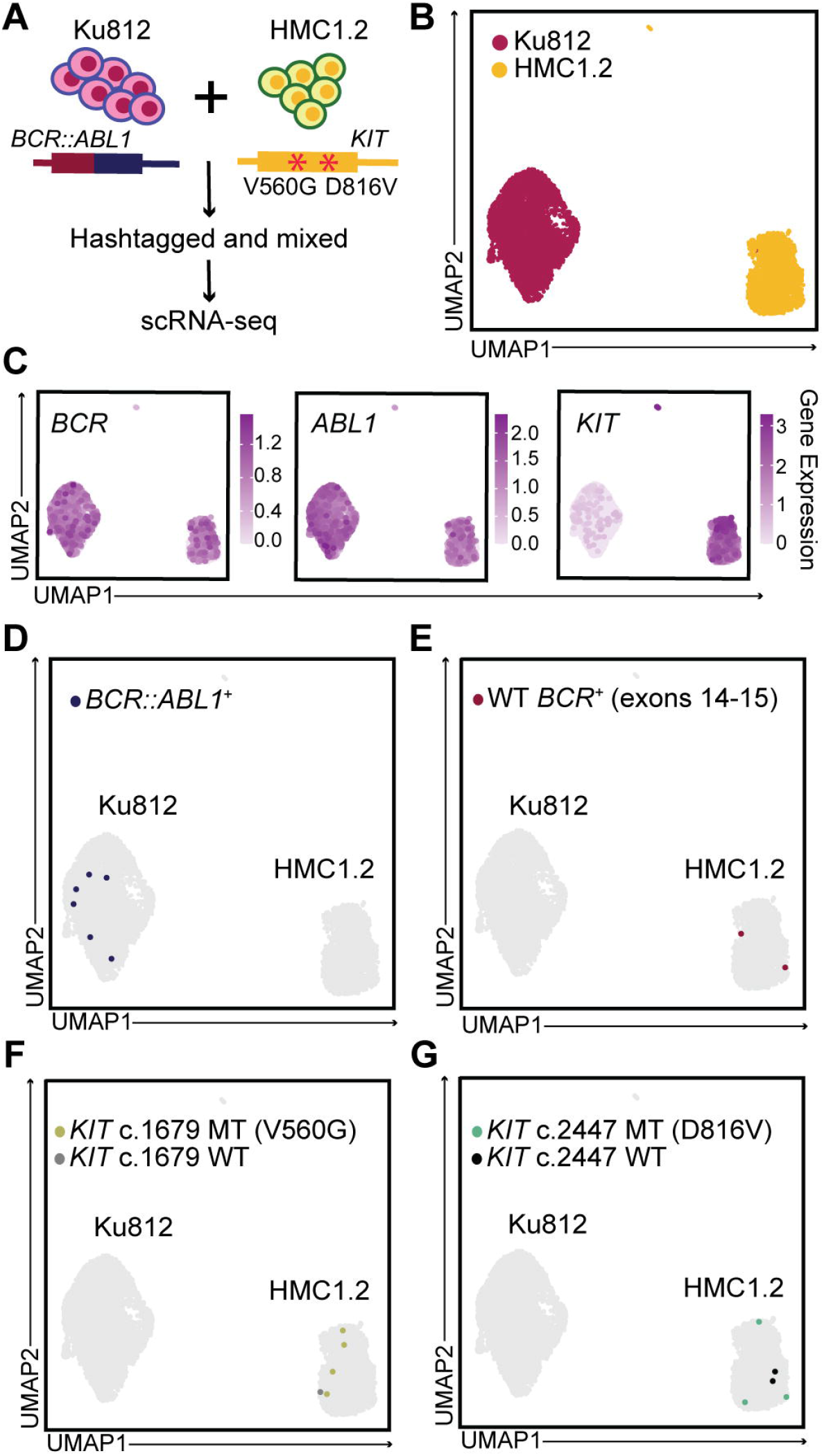
A mixed leukemic cell line system to assess mutation detection in scRNA-seq data. **(A)** Overview of the experimental design. **(B)** UMAP visualization of the scRNA-seq data. The colors represent the cell line assignment of each cell based on hashtag oligo reads. **(C)** UMAP visualizations showing normalized expression of genes of interest. **(D-E)** UMAP visualizations highlighting cells with at least one Illumina read spanning the *BCR::ABL1* fusion breakpoint (D) or the corresponding wild-type *BCR* exon-exon junction (E). **(F-G)** UMAP visualizations highlighting cells with at least one Illumina read containing the indicated *KIT* point mutations or the corresponding wild-type sequence. MT, mutant; WT, wild-type.

As expected, the Ku812 and HMC1.2 cells were transcriptionally distinct (Figure 1B). Plotting the expression of *BCR*, *ABL1*, and *KIT*, irrespective of mutation status, confirmed that the leukemia-associated genes were detected at the single-cell level. *BCR* and *ABL1* expression was observed in both cell lines, whereas *KIT* was mainly expressed in HMC1.2 cells (Figure 1C, Supplementary Figure E1B). To detect *BCR::ABL1* fusion transcripts, we bioinformatically identified the sequencing reads that spanned the *BCR::ABL1* breakpoint. Notably, we detected the *BCR::ABL1* fusion transcript in Ku812 cells but not in HMC1.2 cells, indicating high specificity (Figure 1D). However, only 6 Ku812 cells (0.14 %) were positive, indicating poor sensitivity. Wild-type *BCR* was detected in 2 HMC1.2 (0.08 %), but not in Ku812 cells, consistent with the absence of detectable wild-type *BCR* expression in the Ku812 cells (Figure 1E, Supplementary Figure E1C) [14].

We next interrogated *KIT* transcripts to detect the c.1679T>G (V560G) and c.2447A>T (D816V) mutations, respectively. *KIT* c.1679T>G was detected in 4 HMC1.2 cells (0.15 %), whereas *KIT* c.2447A>T was detected in 3 HMC1.2 cells (0.12 %) (Figure 1F, G). The corresponding wild-type alleles were also detected in HMC1.2, consistent with the heterozygote nature of the mutations. Neither wild-type nor mutant alleles were detected at either *KIT* mutation site in Ku812 cells, in agreement with the virtually undetectable *KIT* expression in these cells.

Taken together, our observations show that the 10X Genomics workflow captures the single-cell transcriptomes of Ku812 and HMC1.2 cells. However, the *BCR::ABL1* fusion transcript and the leukemic point mutations in *KIT* are poorly detected.

### scRNA-seq combined with long-read sequencing enables the detection of leukemic point mutations

The 3’ 10X Genomics single-cell RNA-sequencing workflow captures transcripts using poly(dT) oligos. As a result of cDNA fragmentation during the Illumina library preparation, the sequencing data are inherently biased toward short reads at the 3’ end of the genes. To improve the detection of the mutations of interest, which are located distal to the 3’ end of the transcripts, we developed an alternative long-read PacBio sequencing-based approach. Long-read sequencing analysis of the full-length amplified cDNA that reflects all expressed genes is not cost-effective to detect specific mutations in a few genes. We therefore used a targeted long-read sequencing-based approach to complement the existing scRNA-seq dataset with genotyping data (Figure 2A).

**Figure 2.**
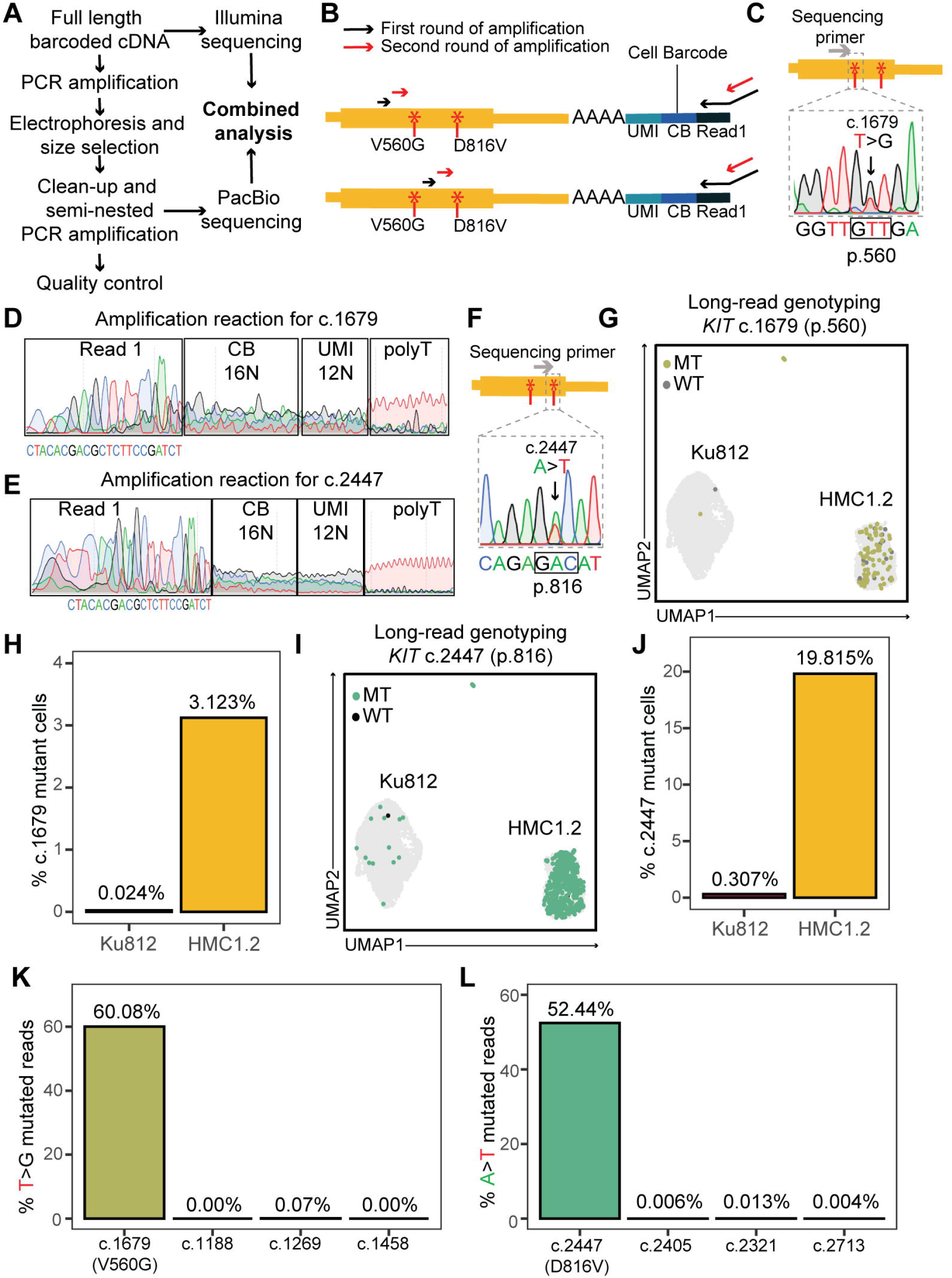
Long-read sequencing enables detection of *KIT* mutations at the single-cell level in HMC1.2 cells. **(A)** Experimental design. **(B)** Schematic of the amplification reactions used for targeted amplification of *KIT* transcripts containing the two point mutations of interest. The black arrows represent primers used in the first amplification round, and the red arrows represent primers used in subsequent amplification steps. The point mutations were captured in two separate reactions using different primer sets. **(C)** Sanger sequencing chromatogram of the amplified *KIT* transcripts prior to long-read sequencing, highlighting the detection of the c.1679T>G mutation. **(D-E)** Sanger sequencing chromatograms showing Read 1, cell barcode and UMI sequences, and the polyT region from the amplification reactions for c.1679 (D) and c.2447 (E). **(F)** Sanger sequencing chromatogram of the amplified *KIT* transcripts prior to long-read sequencing, highlighting the detection of the c.2447A>T mutation. **(G)** UMAP visualization highlighting c.1679T>G mutated (MT) cells in green and c.1679 wild-type (WT) cells in grey. The genotyping is based on the thresholds shown in Supplementary Table E7. **(H)** Bar plot showing the percentage of c.1679 mutant cells in each cell line. **(I)** UMAP visualization highlighting c.2447A>T MT cells in green and c.2447 WT cells in black. The genotyping is based on the thresholds shown in Supplementary Table E7. **(J)** Bar plot showing the percentage of c.2447 mutant cells in each cell line. **(K)** Bar plot showing the percentage of mutant reads (T/(T+G)) at the indicated positions. **(L)** Bar plot showing the percentage of mutant reads (A/(A+T)) at the indicated positions.

Specifically, we utilized remaining amplified full-length cDNA, an intermediate product of the 3’ 10X Genomics workflow, to perform a semi-nested PCR. To detect the *KIT* point mutations, we designed primers that captured A) the mutation sites of interest, B) the cell barcode that identifies the cell to which the sequence belongs, and C) the unique molecular identifier (UMI), which allows us to decipher the number of unique transcripts from each cell (Figure 2B, top). Amplicons of 4.1kb were expected following PCR, measuring from the designed primer site to the polyA tail of the transcript. However, shorter fragments were preferentially amplified, indicating that the 10X Genomics poly(dT) capture oligonucleotide had also bound alternative A-rich regions between the c.1679 (p.560) and c.2447 (p.816) sites of the *KIT* transcript (Supplementary Figure E2A, B and C). This observation prompted us to prepare specific amplicons for each mutation site of interest (Figure 2B).

To ensure that the generated amplicons successfully captured the sites of interest, Sanger sequencing was performed before PacBio sequencing. Sanger sequencing analysis showed the successful capture of the c.1679T>G mutation (Figure 2C) when we sequenced the 5’ end of the associated transcripts. Analysis of the 3’ end detected Read 1, a region corresponding to the cell barcode and UMI, as well as a polyT region indicative of the captured polyA (Figure 2D). The noisy cell barcode and UMI region reflected the generation of a diverse pool of amplicons – a desirable property for the input material for library preparation. These results highlighted the possibility of using Sanger sequencing as a quality control step before long-read sequencing in subsequent experiments. Sanger sequencing of c.2447-associated transcripts revealed the A>T mutation and verified a diverse pool of cell barcodes and UMIs (Figures 2E, F).

We next performed PacBio long-read sequencing of the generated amplicons and integrated the resulting data with the single-cell transcriptomics dataset. The resulting reads containing c.1679 and c.2447 were approximately 1kb in length (Supplementary Figure E2D). Although Ku812 cells are wild-type at c.1679 and c.2447 [15], we detected mutated transcripts among these cells. The detection of these mutant reads likely reflected the 10X Genomics platform’s inherent ability to capture ambient RNA along with the RNA from the cell of interest [16].

To balance sensitivity and false positive rate, we calculated performance metrics across different thresholds of mutation-containing reads per cell (Table 1). Based on this analysis, we established thresholds for labeling a cell as mutant, independent of wild-type allele detection: at least 2 reads for *KIT* c.1679T>G and 4 reads for *KIT* c.2447A>T (Supplementary Figure E2E, F). Applying these thresholds, 3 % of the HMC1.2 cells were considered mutant at c.1679, whereas 0.024 % of the Ku812 cells classified as mutant, reflecting the sensitivity and false positive rate, respectively (Figure 2G, H). For c.2447, 19.8 % of all HMC1.2 cells scored positive for the mutation, reflecting relatively high sensitivity (Figure 2I, J). Only 0.307 % of the Ku812 cells were assigned mutated for c.2447, indicating a low false positive rate (Figure 2J). Notably, the positive predictive value was close to 1 for each mutation (99 % and 98 % for c.1679 and c.2447, respectively). For both mutation sites, genotyped cells showed higher *KIT* expression than non-genotyped cells (Supplementary Figure E2G), supporting an association between target transcript abundance and genotype recovery.

**Table 1.**
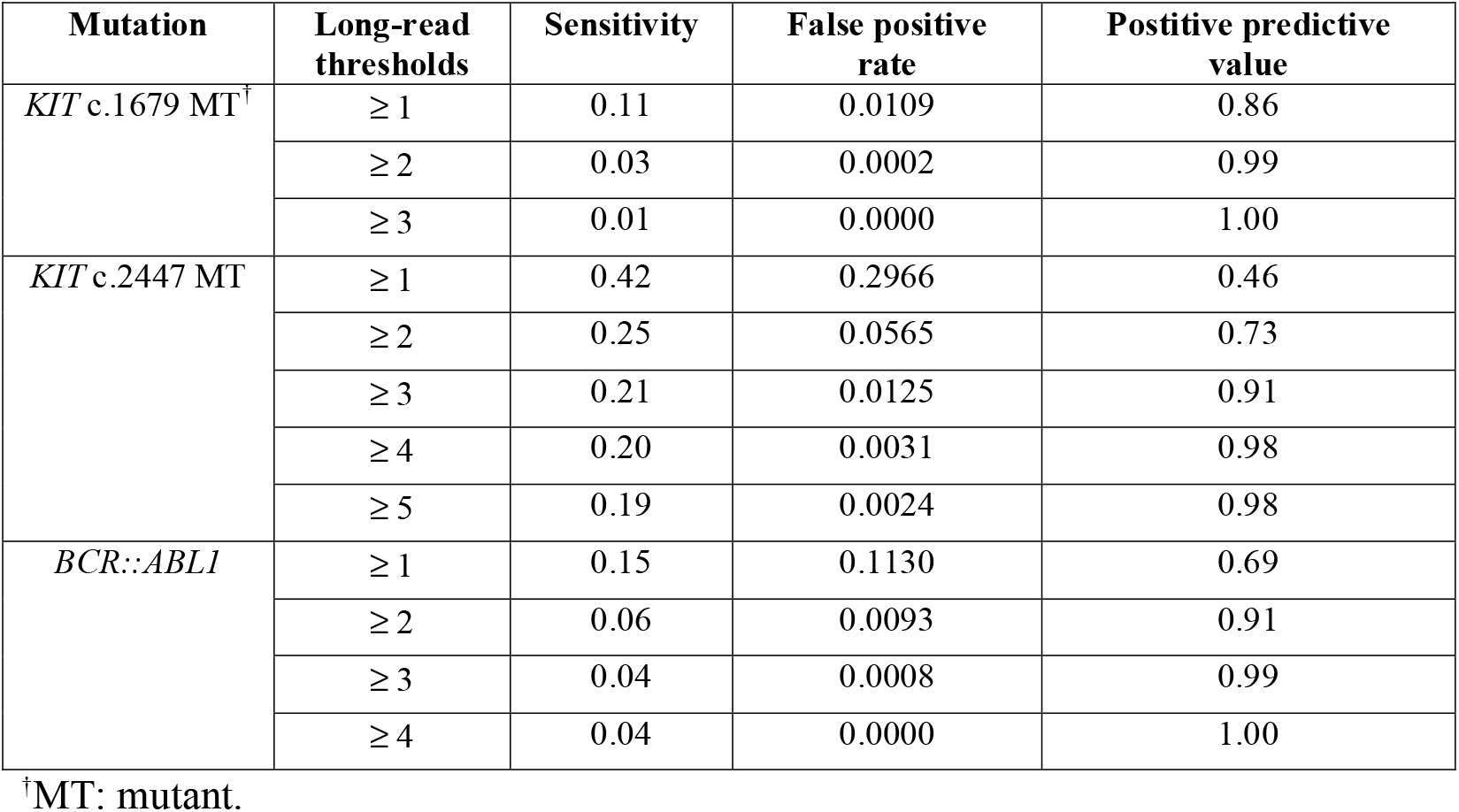
Performance metrics for mutant cell identification at different long-read count thresholds.

We detected the T>G mutation in 60 % of the reads associated with the analysis of *KIT* c.1679 and A>T mutation in 52 % associated with the analysis of c.2447 (Figure 2K, L). To quantify the frequency of spurious mutations, e.g. resulting from PCR and sequencing artifacts, we analyzed multiple positions in *KIT* that were expected to be wild-type. We selected c.1188, c.1269, and c.1458 to quantify spurious T>G mutations. No substitutions were detected at c.1188 and c.1458, whereas 0.07 % of the reads that detected c.1269 were associated with a T>G substitution (Figure 2K). Quantifying the frequency of spurious A>T events revealed up to 0.013 % mutated reads (Figure 2L). Alternative nucleotide substitutions at the mutation sites and control loci were detected only at low frequencies (Supplementary Figure E2H-I). These observations highlight that spurious mutations in *KIT* are negligible compared with the mutations of interest.

Taken together, integration of the 10X Genomics workflow with targeted long-read sequencing allows simultaneous analysis of transcriptomes and point mutations at the single-cell level.

### Long-read sequencing enables simultaneous capture of BCR::ABL1 fusion transcripts and transcriptomes

Chronic myeloid leukemia is associated with the *BCR::ABL1* fusion gene. To detect *BCR::ABL1* fusion transcripts along with the transcriptome of individual Ku812 cells, we designed primers in *BCR* upstream the *BCR::ABL1* breakpoint (Figure 3A). We detected amplicons that were shorter than the expected product (Supplementary Figure E3A, B), as we did for the targeted amplification of *KIT*. Sanger sequencing analysis of the 5’ end of the resulting amplicons revealed a clean *BCR* sequence up to the expected breakpoint and a mixture of bases following the expected fusion site (Figure 3B). Deconvoluting the sequence revealed a superposition of *ABL1* from the fusion transcript and wild-type *BCR*. Because this amplified material was generated from the mixed cell line sample, this signal is consistent with simultaneous amplification of Ku812-derived *BCR::ABL1* transcripts and HMC1.2- derived wild-type *BCR* transcripts. Sanger sequencing of the 3’ end detected Read 1, a diversity of cell barcodes and UMIs, and the polyT region corresponding to the captured polyA (Supplementary Figure E3C).

**Figure 3.**
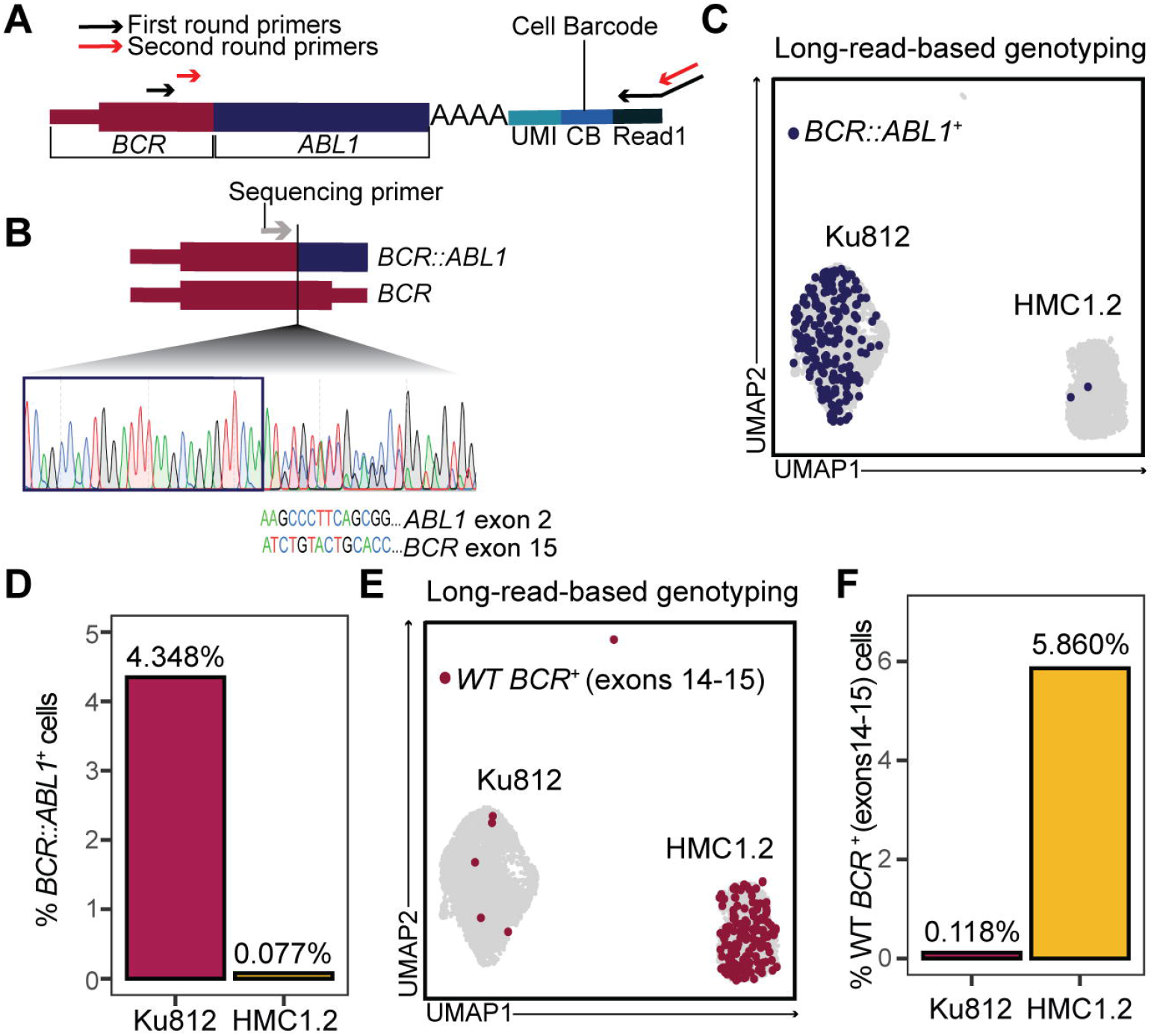
Long-read sequencing enables detection of the *BCR::ABL1* fusion at the single-cell level in Ku812 cells. **(A)** Schematic of the amplification reaction to capture *BCR::ABL1* and wild-type *BCR* transcripts. The black arrows represent primers used in the first amplification round, and the red arrows represent primers used in subsequent amplification steps. Both *BCR::ABL1* and wild-type *BCR* are amplified in the same reaction using a shared set of primers. **(B)** Sanger sequencing chromatogram of the amplified material, highlighting the detection of both the fusion and the wild-type *BCR* in bulk prior to the long- read sequencing, as shown by the mixed signal observed beyond the breakpoint. The sequencing primer used is shown in grey. The box marks exon 14 of *BCR*. **(C)** UMAP visualization highlighting cells in which the *BCR::ABL1* fusion is detected in the long-read sequencing data. The genotyping is based on the thresholds shown in Supplementary Table E7. **(D)** Bar plot showing the percentage of *BCR::ABL1*^+^ cells in each cell line. **(E)** UMAP visualization highlighting cells in which wild-type (WT) *BCR* was detected in the long-read sequencing data. The genotyping is based on the thresholds shown in Supplementary Table E7. **(F)** Bar plot showing the percentage of WT *BCR^+^* cells in each cell line.

The amplicons were subjected to long-read sequencing. The resulting *BCR* or *BCR::ABL1* reads were approximately 2kb in length (Supplementary Figure E3D). As observed for the *KIT* c.2447A>T mutation, *BCR::ABL1* transcripts were detected in both Ku812 and HMC1.2 cells, reflecting the unintentional analysis of ambient RNA. We established a threshold of at least 3 *BCR::ABL1* counts to label a cell as mutant (Table 1, Supplementary Figure E3E).

Applying this threshold, 4.3 % of all Ku812 cells were classified as *BCR::ABL1*^+^, reflecting the sensitivity of the method (Figure 3C, D). At the same threshold, only 0.077 % of the HMC1.2 cells were genotyped as mutant, representing a low false positive rate (Figure 3C, D). Thus, the positive predictive value of the *BCR::ABL1* genotyping reached 99 %.

Cells were classified as wild-type (non-rearranged) *BCR*-positive if they had at least 3 wild-type *BCR* reads in the absence of *BCR::ABL1* reads (Supplementary Table E7). This analysis classified 5.86 % of the HMC1.2 cells and 0.12 % of the Ku812 cells as wild-type *BCR*- positive (Figure 3E, F and Supplementary Figure E3F), consistent with the expected absence of *BCR::ABL1* and detection of wild-type *BCR* in HMC1.2 cells (Supplementary Figure E1C).

Cells genotyped as *BCR::ABL^+^* or wild-type *BCR^+^* showed higher *BCR* expression than non-genotyped cells (Supplementary Figure E3G). In addition, *BCR::ABL1^+^* cells showed higher expression of *ABL1* expression than cells not genotyped as *BCR::ABL1^+^* (Supplementary Figure E3H). These observations further support an association between target transcript abundance and genotype recovery.

Taken together, single-cell RNA-sequencing combined with targeted long-read sequencing enabled detection of *BCR::ABL1*-positive and wild-type/non-rearranged *BCR*-positive cells while allowing analysis of their transcriptomes.

### Integrating scRNA-seq and long-read sequencing captures the transcriptome and disease-driving mutation in chronic myeloid leukemia

To evaluate whether integrated scRNA-seq and long-read sequencing captures *BCR::ABL1* in primary cells, we analyzed leukocytes from a patient with chronic myeloid leukemia (Figure 4). Samples were collected and analyzed at diagnosis and during imatinib treatment to follow the decline in *BCR::ABL1^+^* cells. The 10X Genomics single-cell RNA-seq workflow captured the transcriptomes of lymphocytes, monocytes, and granulocytes (Figure 4A-F). To explore the potential in detecting *BCR::ABL1* transcripts, we plotted the gene expression of *BCR* and *ABL1* based on the Illumina data. Reads associated with each gene were detected at diagnosis and during treatment, prompting a detailed investigation into whether *BCR::ABL1* fusion transcripts can be detected (Figure 4C, F and Supplementary Figure E4A).

**Figure 4.**
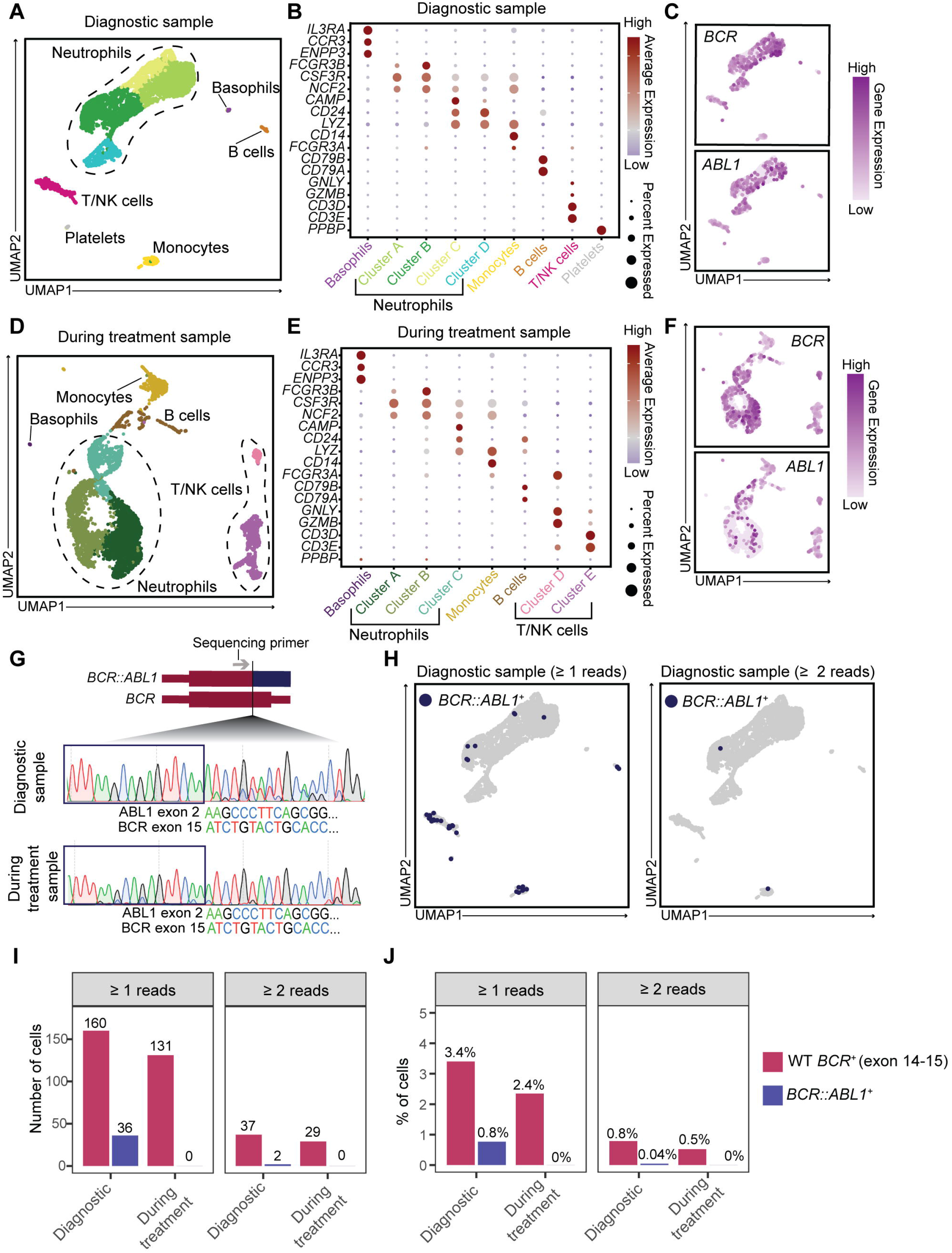
Single-cell detection of *BCR::ABL1* in chronic myeloid leukemia at diagnosis but not during treatment. **(A)** UMAP visualization of 4703 leukocytes from the diagnostic sample, colored by cluster. **(B)** Dot plot of marker genes used to annotate the clusters shown in panel A. **(C)** UMAP visualizations showing normalized expression of selected genes in the diagnostic sample. **(D)** UMAP visualization of 5576 leukocytes from the sample collected during treatment, colored by cluster. **(E)** Dot plot of marker genes used to annotate the clusters shown in panel D. **(F)** UMAP visualizations showing normalized expression of selected genes in the sample collected during treatment. **(G)** Sanger sequencing chromatograms of the amplified material at diagnosis (top) and during treatment (bottom). The Sanger sequencing primer is shown in grey. The boxes mark exon 14 of *BCR*. **(H)** UMAP visualization of leukocytes from the diagnostic sample highlighting *BCR::ABL1^+^* cells identified at different long-read thresholds. **(I)** Number of cells with at least 1 (left) or at least 2 (right) long-read sequencing reads for wild-type (WT) *BCR* (pink) or *BCR::ABL1* (purple). **(J)** Percentage of cells identified as wild-type *BCR^+^* or *BCR::ABL1^+^* at the indicated thresholds.

We performed semi-nested PCR to amplify the *BCR::ABL1* and/or *BCR* transcripts from the 10X Genomics-derived full-length cDNA of each sample. Sanger sequencing of the diagnostic sample showed a clear *BCR* signature up to the prospective breakpoint, from which the sequences were mixed and reflected a superposition of *BCR* and *ABL1* signals (Figure 4G). The sample obtained during treatment did not show a clear *ABL1* signal downstream the breakpoint, indicating reduced levels of fusion transcripts (Figure 4G). As in the cell line experiment, we detected Read 1, diverse cell barcodes and UMIs, and the polyT sequence (Supplementary Figure E4B).

Long-read sequencing analysis of the diagnostic sample detected the wild-type *BCR* as well as *BCR::ABL1* fusion transcripts (Supplementary Figure E4C, D). Unlike the cell line genotyping, establishing a threshold count to confidently annotate primary patient cells as mutant is challenging. We therefore plotted the data using different thresholds (Figure 4H-J, Supplementary Figure E4E). A threshold of 1 sequencing read detected *BCR::ABL1* across the hematopoietic cell types at diagnosis. Increasing the threshold to 2 sequencing reads restricted the expression to a neutrophil and a monocyte, which both showed high expression of *BCR::ABL1* (28 and 329 UMIs respectively). Wild-type *BCR* was detected at the same order of magnitude at diagnosis and during treatment (Figure 4I, J). Strikingly, no cells with detectable *BCR::ABL1* were identified during treatment, irrespective of threshold. This was in line with the reduction of *BCR::ABL1* levels measured in the clinic (41% at diagnosis and 0.1% during treatment). Further analysis revealed that genotyped cells displayed higher expression of the corresponding genes than non-genotyped cells, consistent with the cell line results (Supplementary Figure E4F-H).

Taken together, the integration of scRNA-seq and long-read sequencing analysis enables genotyping of *BCR::ABL1* in primary cells.

## Discussion

Here, we systematically established a method that simultaneously detects leukemic mutations and captures the transcriptomes of single cells. Our approach uses the stable amplified cDNA from the 10X Genomics experimental workflow as input material for the genotyping. Thus, the genotyping for one or more specific mutations can be performed on already processed samples. This does not remove the need to define a target gene or region for primer design, but it allows the decision to be made after the standard 10X Genomics processing has been completed. The GoT pipeline includes mutation-customized primers during the first cDNA amplification step, which requires the mutations of interest to be characterized before or immediately upon initiating the 10X Genomics-based single-cell RNA-sequencing experiment [4]. GoT is therefore inapplicable for many types of diagnostic samples, especially when the cells of interest do not tolerate freeze-thawing cycles and the mutation profile is unknown.

The GoT method was initially tested to be compatible with the Oxford Nanopore long-read sequencing platform [4]. However, GoT is primarily designed for the Illumina short-read platform. The short-read sequencing requires the mutation site of interest, often located distant from the 3’ end, to be brought close to the cell barcode and UMI for sequencing. This is achieved through sequential rounds of volume-sensitive intramolecular circularization steps and inverse PCRs, allowing for the resulting molecules to be analyzed with short-read Illumina sequencing. In contrast, our method applies semi-nested PCR to already amplified cDNA and uses PacBio long-read sequencing to directly read the targeted region together with the 10X Genomics cell barcode and UMI from the same amplicon.

The 10X Genomics-based capture oligos are expected to bind mainly the polyA tail of the transcript. However, our data show that the capture oligos also bind within genes, likely A- rich stretches. This phenomenon is important to be aware of when preparing long-read sequencing libraries, as molecules of shorter length with the mutations of interest might be excluded by a strict size selection. Such secondary priming positions have previously been used to infer RNA velocity [17], and we now show its applicability for genotyping.

Our method uses target-specific semi-nested PCR to enrich transcripts for mutation analysis. Hybridization capture followed by universal PCR provides an alternative strategy for enriching specific transcripts and is used in LOTR-seq [8, 9]. As implemented, our method uses separate PCR reactions, enabling assay optimization and quality control of individual targets before long-read sequencing. LOTR-seq enriches all targets together, enabling parallel analysis of a larger gene panel but limiting separate optimization and quality control of individual targets (see Supplementary Table E8 for a comparison of the two methods).

We successfully genotyped 4.3 % of the *BCR::ABL1*-carrying Ku812 cells using the 3’ platform of 10X Genomics. Similarly, Penter et al genotyped another *BCR::ABL1*-carrying cell line, K562, using the 10X Genomics 5’ platform. In that study, the fusion gene was captured in 2.7 % of the individual K562 cells, using the Oxford Nanopore platform [10]. The fraction of genotyped cells should be interpreted in light of the post-hoc design. Importantly, in our controlled mixed cell line system where ground truth is known, the method enabled high-confidence annotation of mutant cells, with positive predictive values close to 1.

For the point-mutation analysis, we also evaluated whether the detected *KIT* mutations could be explained by technical artifacts introduced during PCR or sequencing. This analysis showed that substitutions at the expected wild-type positions in *KIT* were negligible compared with the mutations of interest. These results suggested that the low-level *KIT* mutant signal in Ku812 cells reflected ambient RNA rather than PCR or sequencing artifacts, underscoring a broader limitation of transcript-based genotyping in droplet-based single-cell RNA-sequencing.

All methods that genotype cells based on RNA have the inherent limitation that only cells that express the genes of interest can be confidently genotyped. RNA-based genotyping methods therefore have the potential to chart the distribution of the somatic *KIT* c.2447A>T (D816V) and c.1679T>G (V560G) mutations in KIT-expressing hematopoietic progenitor cells and mature mast cells, but not necessarily in other mature hematopoietic cell types where KIT expression is lacking. On the other hand, activating mutations such as KIT D816V are mainly expected to dysregulate cells that express the gene and protein.

Our cell line mixing experiment, in which we used cell type-specific hashtags to identify each cell, ensured that we could closely monitor the performance of the genotyping. Expression of the gene of interest is, as already discussed, critical to genotype the cells. Beyond this, the controlled setup revealed a broader interpretive issue for transcript-based single-cell genotyping: the detection of a mutant transcript is not necessarily proof that a given cell is mutant. For this reason, requiring more mutation-containing reads makes a mutant annotation more confident, but also increases the risk of missing mutant cells. Cell line mixtures provide a controlled setting for threshold selection, because the expected mutation status of each cell line is known. For primary cells, it is a significant challenge to set the threshold, as the ground truth is unknown and the background noise will depend on several parameters. Firstly, highly expressed genes are expected to contribute more to the ambient RNA pool than lowly expressed genes, and secondly deep sequencing of diverse libraries is likely to pick up ambient RNA to a higher extent than shallow sequencing of low-complexity libraries. Different thresholds can therefore be evaluated depending on how much confidence is required when assigning mutation status to individual cells. A universal threshold for genotyping can therefore not be applied, and cells lacking informative target transcripts should remain unassigned rather than being annotated as mutant or wild-type. This is particularly important for comparisons between mutant and wild-type cells, which require both groups to be confidently annotated.

Although our RNA-based genotyping can be applied to already processed samples, the targeted semi-nested PCR amplification requires the potential mutation site of interest to be determined. The HMC1.2 and Ku812 cells that we analyzed harbor well-characterized mutations in *KIT* and *BCR::ABL1*, respectively. For the patient material, we performed Sanger sequencing of bulk cells to identify the disease-driving *BCR::ABL1* transcript before the single-cell genotyping. The long-read PacBio sequencing can in theory be used to identify and characterize additional mutations in proximity to the regions of interest if they fall within the amplified fraction. However, near-full length transcripts were challenging to capture, likely due to A-rich regions of the RNA binding the capture oligos.

Taken together, we developed and systematically evaluated a targeted long-read sequencing approach for post-hoc genotyping of cells profiled with the 10X Genomics 3’ workflow. By linking leukemia-associated mutation-containing transcripts to transcriptionally defined cells, the approach extends the information that can be recovered from existing single-cell RNA-sequencing experiments. Our findings also provide guidance for interpreting transcript-based genotype assignments in single-cell data.

## Authors contributions

S.P., C.W., D.B., U.O-S. and S.S. and J.S.D. conceptualized the study. S.P., C.W., L.M., and J.M. performed the experiments. U.O-S. and S.S. coordinated the collection of primary samples. S.P. and D.B. performed the bioinformatics analysis. S.P., D.B., C.W., L.M., J.M., and J.S.D. contributed to the data analysis and interpretation. S.P., C.W., J.M., G.N. and J.S.D. designed the experiments. S.P. and J.S.D. drafted the manuscript. J.S.D. supervised the study and secured funding.

## Data availability

Single-cell RNA-sequencing data have been deposited in the Gene Expression Omnibus under accession number GSE329598. Raw Illumina and PacBio long-read sequencing data generated from patient samples are not publicly available, as these data are considered sensitive personal data under Swedish legislation and the General Data Protection Regulation. Access to these raw data may be requested from the corresponding author and is subject to fulfillment of all applicable legal and ethical requirements.

## Funding statement and acknowledgements

The study was supported by grants from the Swedish Research Council (2022-00558 and 2025-00352), the Swedish Cancer Society, the Cancer Research Foundations of Radiumhemmet, Karolinska Institutet, and Magnus Bergvall’s Foundation. S.P. was supported by a scholarship from the Onassis Foundation (scholarship ID: F ZO 057-2/2022-2023). Related to Illumina sequencing: the authors acknowledge support from the National Genomics Infrastructure in Stockholm funded by Science for Life Laboratory, the Knut and Alice Wallenberg Foundation and the Swedish Research Council, and NAISS for assistance with massively parallel sequencing and access to the UPPMAX computational infrastructure. Related to long-read sequencing: the authors would like to acknowledge support of the National Genomics Infrastructure (NGI) / Uppsala Genome Center and UPPMAX for providing assistance in massive parallel sequencing and computational infrastructure. Work performed at NGI / Uppsala Genome Center has been funded by RFI / VR and Science for Life Laboratory, Sweden

## Conflicts of interest statement

Chenyan Wu is currently employed by Genoskin. Genoskin had no role in the study. All other authors declare no conflicts of interest.

## Ethics approval statement

The Swedish Ethical Review Authority approved the study (2022-02570-01)

## Patient consent statement

Patient samples were collected after written informed consent.

## Supporting information

Supplementary Script

## Notes

### Summary of Updates

Updated title; updated figures; updated supplement; added quality control metrics, refocused the contents.

