## Supplementary figures and images for "Post-hoc long-read sequencing links leukemic mutation status to single-cell transcriptomes"

### Supplementary Figure E1

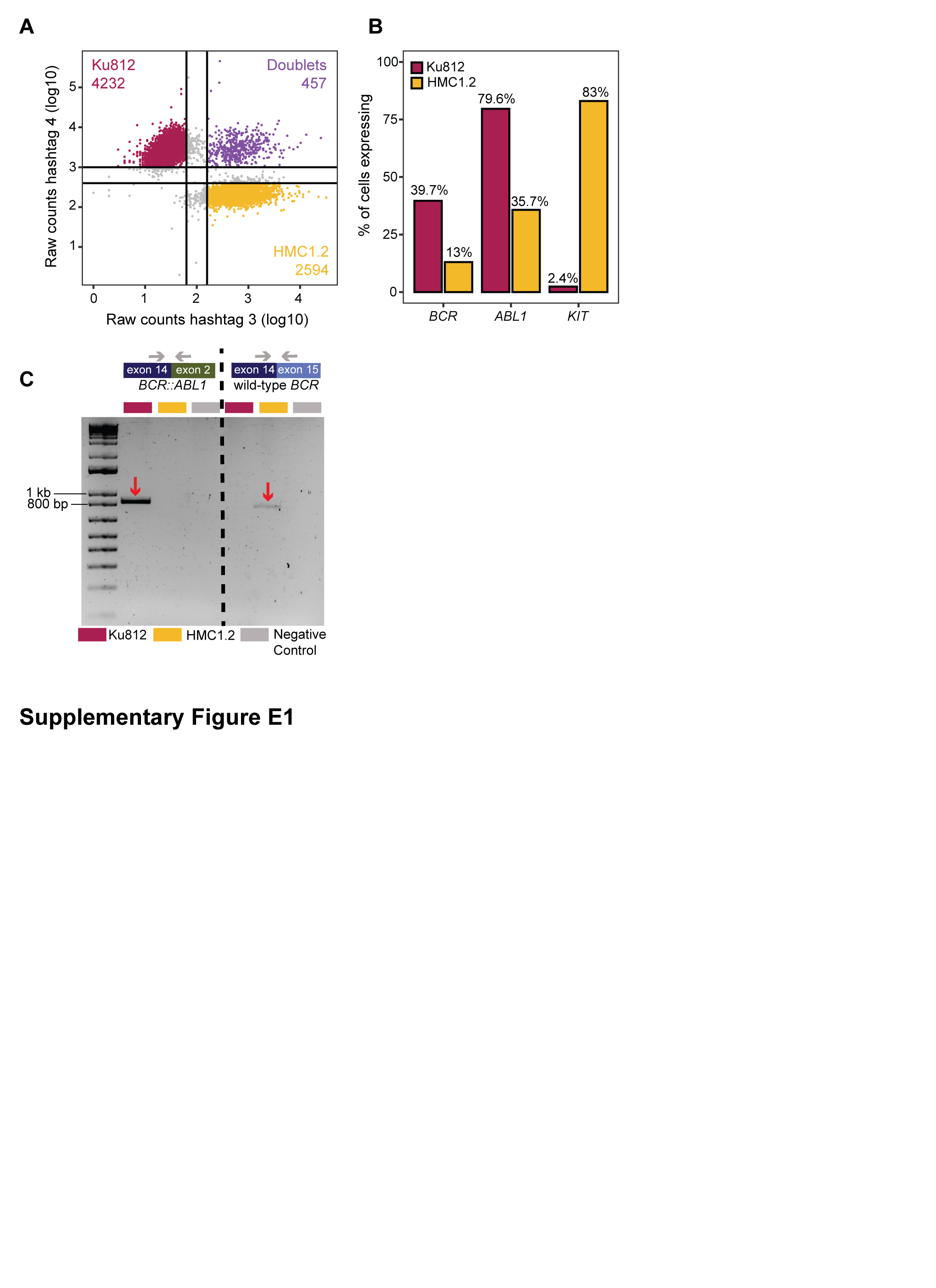

### Supplementary Figure E2

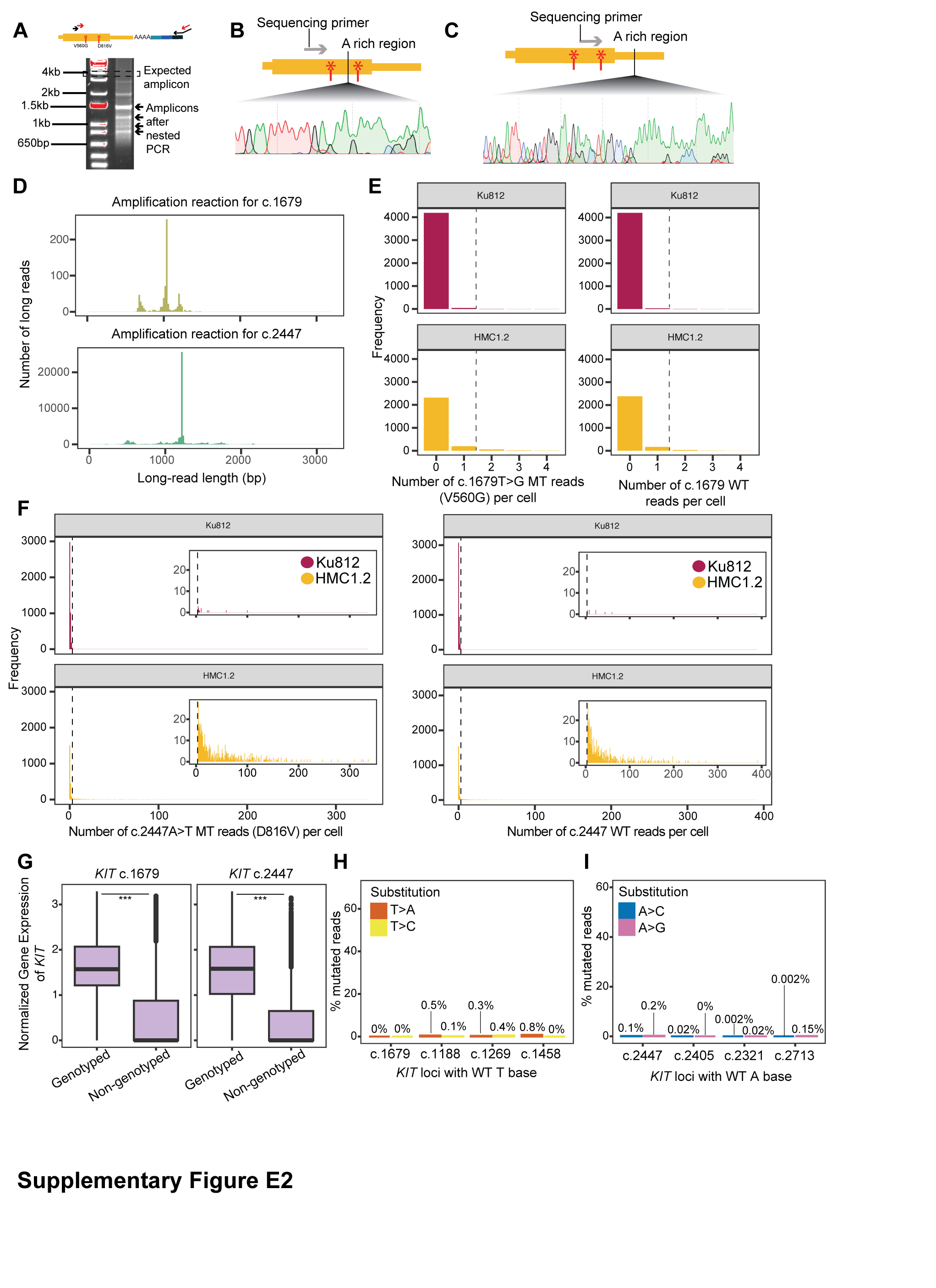

### Supplementary Figure E3

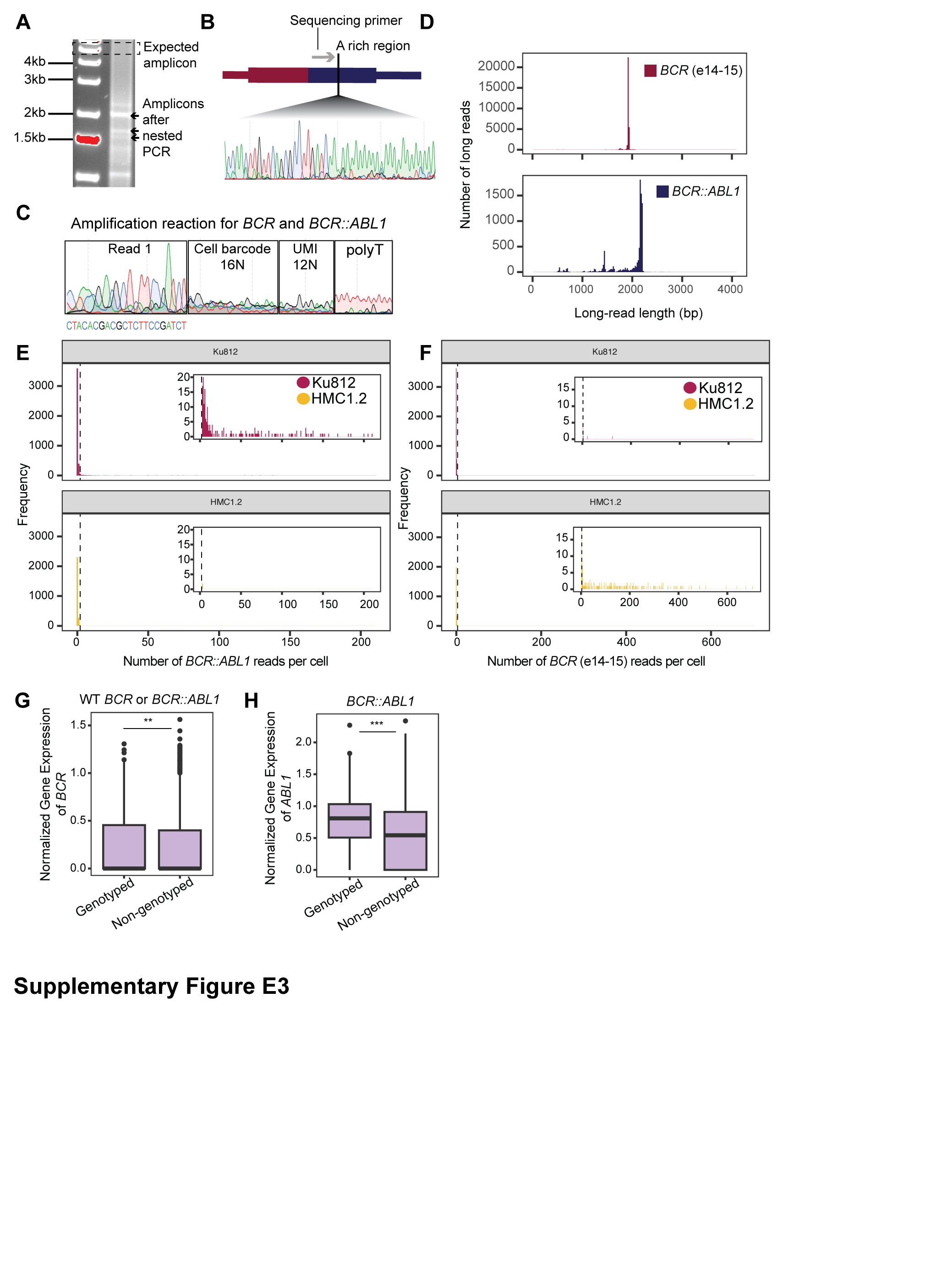

### Supplementary Figure E4

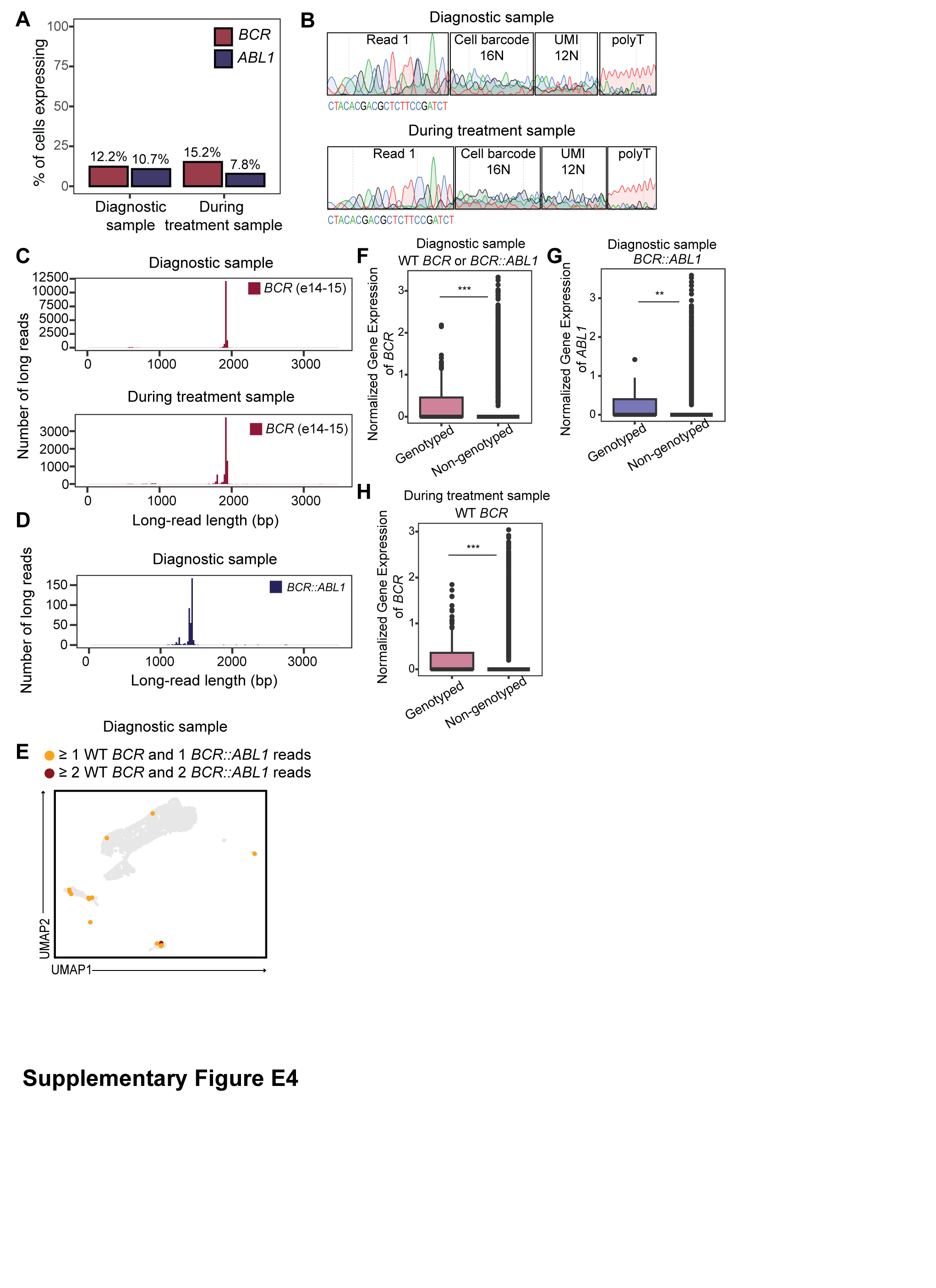
