## Supplementary Script for "Post-hoc long-read sequencing links leukemic mutation status to single-cell transcriptomes"

# Post-hoc long-read sequencing links leukemic mutation status to single-cell transcriptomes
# Sofia Papavasileiou,1 Chenyan Wu,1 Daryl Boey,1 Lucille Margerie,1 Jiezhen Mo,1 Ulla Olsson-Strömberg,2 Stina Söderlund,2 Gunnar Nilsson,1,2 and Joakim S. Dahlin1*

### This supplementary script is a template for pre-processing of the long-read PacBio genotyping data.

######### Step 1: Identify long reads with unique occurrence of Read 1. #########

###### Separate the forward strand reads from the reverse strand reads.
#### The fastq file has reads in both strands and I am separating them using read1 sequence.
### All the reads will have on the left side the Read 1 followed by cell barcode and uni (referred to as forward reads) so the gene specific sequences will be in reverse.

# Keep reads that the read 1 was identified only once.
gzcat <Input_fastq_file.fastq.gz> | grep "CTACACGACGCTCTTCCGATCT" | grep -v "CTACACGACGCTCTTCCGATCT.*CTACACGACGCTCTTCCGATCT" |grep -v "AGATCGGAAGAGCGTCGTGTAG" > forward.fastq


# Keep reads that the reverse sequence of read 1 was identified only once in a file
gzcat <Input_fastq_file.fastq.gz> | grep "AGATCGGAAGAGCGTCGTGTAG" | grep -v "AGATCGGAAGAGCGTCGTGTAG.*AGATCGGAAGAGCGTCGTGTAG" | grep -v "CTACACGACGCTCTTCCGATCT" > reverse.fastq


# Get the reverse complement of the reversed reads
cat reverse.fastq | awk '{print $1}' | tr 'ATCGatcg' 'TAGCtagc' | rev > reverse_corrected.fastq


# Add the files that have been reverse complemented together with the forward in one file excluding the ones that read 1 was not detected or detected twice
cat forward.fastq reverse.fastq > cleaned.fastq


#### Identify read1 in all the reads and remove everything that comes before that for all of them to be aligned.
awk '{print $1}' cleaned.fastq | sed 's/^.*CTACACGACGCTCTTCCGATCT/CTACACGACGCTCTTCCGATCT/' > cleaned_aligned.fastq

######### Step 2: Extract Cell barcode and UMI. #########

#### The bases directly after Read1 correspond to the cell barcode and UMI. 
### The sequences will be extracted and removed from the read.

# Keep the read1 in a file
awk '{print $1}' cleaned_aligned.fastq | cut -c-22 > Read1.txt

# Remove read 1 from the sequence. I need to add the position I need to keep which is one more than the bases I trim
cat cleaned_aligned.fastq | cut -c 23- > sequences.txt 

# Keep cell barcode in a file
awk '{print $1}' sequences.txt | cut -c-16 > Cell_barcodes.txt

# Remove cell barcode from the sequence
cat sequences.txt | cut -c 17- > sequences1.txt

# Keep UMI in a file
awk '{print $1}' sequences1.txt | cut -c-12 > UMI.txt

# Remove UMI from the sequence
cat sequences1.txt | cut -c 13- > sequences2.txt

# Making an ID based on the cell barcode and the UMI
paste -d "\0" Cell_barcodes.txt UMI.txt > IDs.txt

# Keep a column with the read length
cat sequences2.txt | awk '{print length}' > read_length.txt


# Putting everything together
paste Read1.txt Cell_barcodes.txt UMI.txt IDs.txt sequences2.txt read_length.txt > Reads_table.txt

######### Step 3: Isolate only the reads that have one of the forward primers detected. #########

## BCR/BCR::ABL1 primer
cat Reads_table.txt | grep "GATCTTGATCTTCAAAGCGTCCA\>" | grep -v "GATCTTGATCTTCAAAGCGTCCA.*GATCTTGATCTTCAAAGCGTCCA" | grep -v "CTCGTCATCCTCCATGATGGC" | grep -v "GATCCACTGCTGGTGTTC" > PrimerA_reads.txt
cat PrimerA_reads.txt | awk '{print $1,$2,$3,$4,$5,$6,"PrimerA"}' > PrimerA_reads_cleaned.txt

## c.2447 (D816V) primer (note this refers to primer C2 in Supplementary Table E2)
cat Reads_table.txt | grep "CTCGTCATCCTCCATGATGGC\>" | grep -v "CTCGTCATCCTCCATGATGGC.*CTCGTCATCCTCCATGATGGC" | grep -v "GATCTTGATCTTCAAAGCGTCCA" | grep -v "GATCCACTGCTGGTGTTC" > PrimerB_reads.txt
cat PrimerB_reads.txt | awk '{print $1,$2,$3,$4,$5,$6,"PrimerB"}' > PrimerB_reads_cleaned.txt

## c.1679 (V560G) primer (note this refers to primer B2 in Supplementary Table E2)
cat Reads_table.txt | grep "GATCCACTGCTGGTGTTC\>" | grep -v "GATCCACTGCTGGTGTTC.*GATCCACTGCTGGTGTTC" | grep -v "GATCTTGATCTTCAAAGCGTCCA" > PrimerC_reads.txt
cat PrimerC_reads.txt | awk '{print $1,$2,$3,$4,$5,$6,"PrimerC"}' > PrimerC_reads_cleaned.txt

######### Step 4: Remove PCR duplicates in files #########
## BCR primers
cat PrimerA_reads_cleaned.txt | sort -u -k4,4 > PrimerA_reads_deduplicated.txt

## c.2447 (D816V) primer
cat PrimerB_reads_cleaned.txt | sort -u -k4,4 > PrimerB_reads_deduplicated.txt

## c.1679 (V560G) primer
cat PrimerC_reads_cleaned.txt | sort -u -k4,4 > PrimerC_reads_deduplicated.txt


######### Step 5: Deconvolute BCR and BCR::ABL1. #########
### The KIT MT and WT read deconvolution will happen at a later stage in R.

cat PrimerA_reads_deduplicated.txt | grep "AGGGTGCAGTACAGATTTGAACTCTGCTTAAA" | grep -v "AGGGTGCAGTACAGATTTGAACTCTGCTTAAA.*AGGGTGCAGTACAGATTTGAACTCTGCTTAAA" | grep -v "GCCGCTGAAGGGCTTTTGAACTCTGCTTAAAT" > BCR_reads.txt
cat PrimerA_reads_deduplicated.txt | grep "GCCGCTGAAGGGCTTTTGAACTCTGCTTAAAT" | grep -v "GCCGCTGAAGGGCTTTTGAACTCTGCTTAAAT.*GCCGCTGAAGGGCTTTTGAACTCTGCTTAAAT" | grep -v "AGGGTGCAGTACAGATTTGAACTCTGCTTAAA" > BCR_ABL1_reads.txt


cat BCR_reads.txt | awk '{print $1,$2,$3,$4,$5,$6,$7,"BCR"}' > BCR_reads_annotated.txt
cat BCR_ABL1_reads.txt | awk '{print $1,$2,$3,$4,$5,$6,$7,"BCR-ABL1"}' > BCR_ABL1_reads_annotated.txt

cat PrimerB_reads_deduplicated.txt | awk '{print $1,$2,$3,$4,$5,$6,$7,"KIT"}' > KIT_D816V_reads.txt
cat PrimerC_reads_deduplicated.txt | awk '{print $1,$2,$3,$4,$5,$6,$7,"KIT"}' > KIT_V560G_reads.txt


cat BCR_reads_annotated.txt BCR_ABL1_reads_annotated.txt KIT_D816V_reads.txt KIT_V560G_reads.txt > Cell_lines_Filtered_long_reads.txt
