## Supplementary Appendix for "Post-hoc long-read sequencing links leukemic mutation status to single-cell transcriptomes"

### Supplementary Methods

*Cell line culture*

The Ku812 cells were cultured in RPMI medium (Cytiva HyClone, Global Life Sciences Solutions, Marlborough, MA, USA) supplemented with 2 mM L-glutamine (Cytiva HyClone), 10% heat-inactivated fetal bovine serum (Gibco / Thermo Fisher Scientific, Waltham, MA, USA), and 100 U/mL penicillin and 0.1 mg/mL streptomycin (Cytiva HyClone). The HMC1.2 cells were cultured in IMDM medium (Gibco / Thermo Fisher Scientific) supplemented with 2 mM L-glutamine (Cytiva HyClone), 10% heat-inactivated fetal bovine serum (Gibco / Thermo Fisher Scientific), 100 U/mL penicillin and 0.1 mg/mL streptomycin (Cytiva HyClone), and 65 μL monothioglycerol (Sigma-Aldrich, St. Louis,

Missouri, USA) per 500 mL of medium. Both cell lines were authenticated based on their STR profiles (Eurofins Genomics, Louisville, Kentucky, USA). In addition, Sanger sequencing verified the presence of the *BCR::ABL1* fusion in the Ku812 cells and *KIT* point mutations in the HMC1.2 cells.

*Leukocyte isolation from patient samples*

To generate the transcriptomic datasets of leukocytes, cells were isolated from 0.5 mL of peripheral blood (at diagnosis) or 1 mL of bone marrow aspirate (during treatment), collected in EDTA tubes. Mature red blood cells were lysed using BD Pharm Lyse (BD Biosciences, Franklin Lakes, New Jersey, USA) and the cells were filtered using a 35 µm filter. To avoid interference from EDTA with cDNA synthesis, the cells were washed in PBS supplemented with 2% heat-inactivated fetal bovine serum before proceeding to the scRNA-seq workflow.

*Bioinformatics analysis of Illumina data*

The fastq files were processed using Cell Ranger (10X Genomics) (version 7.0.1 for the cell lines, version 7.1.0 for the diagnostic sample and version 8.0.1 for the treatment sample). Reads were aligned using the GRCh38-2020-A reference. The Cell Ranger filtered count table was imported into R (version 4.5.2) and analyzed with Seurat [1] (version 5.4.0) using RStudio (version 2026.01.0+392).

For the analysis of the cell line dataset, the raw hashtag-derived reads were used to assign the cells to one of the two cell lines and to identify doublets containing cells from both cell lines. An undetermined cell population was also defined. The thresholds for assignment are shown in Supplementary Figure E1A. Quality control filtering of the dataset included removal of cells with too few or too many features, excess UMIs, or a high percentage of mitochondrial reads (Supplementary Table E3). Lastly, cells that were identified as doublets and undetermined cells were also removed from the dataset.

For the analysis of the patient datasets, cell barcodes with low feature counts were excluded. Cell doublets were simulated using scDblFinder [2], and predicted doublets were removed from the analysis. Additional filtering excluded cell barcodes with unusually high feature counts or elevated mitochondrial read fractions (Supplementary Table E3). Genes detected in fewer than 5 cells were also excluded.

For all datasets, after the initial quality control steps, the raw gene expression counts were normalized using the NormalizeData function. The 2000 variable features were selected using the FindVariableFeatures function and the vst method. The Leiden algorithm (resolution 0.2) was used to identify cell clusters in the patient datasets.

*Detection of the KIT point mutations, wild-type BCR, and BCR::ABL1 in the short-read sequencing data*

The Illumina fastq files were parsed to isolate reads containing either the *BCR::ABL1* breakpoint (*BCR* exon 14 – *ABL1* exon 2), the wild-type (non-rearranged) *BCR* (exons 14-15), the *KIT* region upstream c.1679, or the *KIT* region upstream c.2447. After isolating those reads, the corresponding cell barcodes and unique molecular identifiers (UMIs) were extracted. The reads were further filtered to include only the reads whose cell barcode was found in the filtered dataset. Reads sharing the same cell barcode and UMI were collapsed. The *KIT* reads were assessed for the presence of the wild-type or mutated allele at positions c.1679 and c.2447. The filtered reads were then added to the Seurat object of the short-read sequencing data.

*Detection of BCR::ABL1 and wild-type BCR in cell lines by reverse transcription PCR*

Total RNA from each cell line was isolated using the Quick RNA Microprep Kit (Zymo Research, Irvine, California, USA) according to the manufacturer's recommendations. cDNA was synthesized from the isolated RNA using ProtoScript II First strand cDNA Synthesis Kit (New England Biolabs, Ipswich, Massachusetts, USA) with oligo(dT) primers. Two separate PCR reactions (one for each target molecule) were performed using the same cDNA input and molecule-specific primers to amplify either *BCR::ABL1* or wild-type *BCR* (Supplementary Figure E1C). Agarose gel electrophoresis was used to assess the presence of each transcript in the cell lines.

*PacBio library preparation and sequencing*

The amplified fragments from the cell line experiment were used as input for the SMRTbell library preparation kit (version 3.0) according to PacBio's Procedure and Checklist (PN 102-359-00 REV 02 SEP 2022) (Pacific Biosciences, California, USA). The SMRTbell library was sequenced on the Revio instrument using the Revio binding kit, the Revio sequencing plate and the Revio SMRT Cell 25M with a movie time of 24 hours.

The amplified material from the patient samples was used for the preparation of two SMRTbell libraries using the SMRTbell prep kit (version 3.0) according to PacBio's Procedure and Checklist (PN 102-359-000 REV 03 DEC 2023). Each library was indexed and pooled according to the protocol and the pool was sequenced on the Revio instrument using the Revio binding kit, the Revio sequencing plate and the Revio SMRT Cell 25M with a movie time of 30 hours.

*Pre-processing of the long-read sequencing data*

The fastq files generated after sequencing were processed using a Bash script (Supplementary Script) to remove reads containing multiple Read1 sequences. Cell barcodes and UMIs were extracted, and only reads containing a unique forward primer (A2, B2, or C2; Supplementary Table E2) were retained for downstream analysis. PCR duplicates were removed by collapsing reads with identical cell barcode and UMI combinations and reads were assigned to either *KIT*, wild-type *BCR,* or *BCR::ABL1* (Supplementary Table E6) before import into R (version 4.5.3). The deduplicated reads were then filtered further so that only reads with cell barcodes present in the short-read Illumina-based dataset were included in the analysis. The mutational status of each *KIT* read was determined by identifying sequences according to Supplementary Table E6. Lastly, a long-read count table was generated, and the long-read counts were saved in the Seurat object.

The genotype status of each cell was determined based on the number of long reads detected (Supplementary Table E7). The thresholds were empirically defined based on the read distributions (Table 1).

*Calculating error rate (related to Figures 2K-L and Supplementary Figures E2H-I)*

To assess the capture of spurious mutations arising from amplification or sequencing errors, we calculated the frequency of all possible nucleotide substitutions at the candidate mutation sites and corresponding control genomic positions proximal to c.1679 and c.2447. Briefly, for each position, we determined the number or reads with the base corresponding to the wild-type (WT) allele and the ones corresponding to the mutant (MT) allele. The percentage of mutated reads in each position was then calculated as MT / (MT + WT). All long reads included in the Seurat object were used in this analysis.

### Supplementary Tables

**Table E1. Amplification conditions.**

| Amplification Reaction (Target molecule, Round) | Primers used (Table E2) | Annealing temperature | Extension Time |
|---|---|---|---|
| *BCR/ BCR::ABL1*, Round 1 | A1 and R1 | 66 °C | 3 min 20 s |
| *BCR/ BCR::ABL1*, Round 2 and 3 | A2 and R2 | 66 °C | 3 min 20 s |
| *KIT* c.1679, Round 1 | B1 and R1 | 65 °C | 2 min 20 s |
| *KIT* c.1679, Round 2 and 3 | B2 and R2 | 64 °C | 1 min |
| *KIT* c.2447, Round 1 | C1 and R1 | 60 °C | 1 min 30 s |
| *KIT* c.2447, Round 2 and 3 | C2 and R2 | 66 °C | 1 min 30 s |

**Table E2. Primer sequences used for targeted amplification.**

| Primer Name | 5' → 3' sequence |
|---|---|
| Universal overhang + Read1 primer (R1) | AATGATACGGCGACCACCGAGATCTACACTCTTTCCCTACACGACGCTC |
| Universal Overhang primer (R2) | AATGATACGGCGACCACCGA |
| *BCR* primer Forward_1 (Primer A1) | GCAAAACGCAGCAGTATGAC |
| *BCR* primer Forward_2 (Primer A2) | TGGACGCTTTGAAGATCAAGATC |
| KIT c.1679 Forward_1 (Primer B1) | GGAGTGTTCATGTGTTATGCC |
| KIT c.1679 Forward_2 (Primer B2) | GAACACCAGCAGTGGATC |
| KIT c.2447 Forward_1 (Primer C1) | CCAACACAACTTCCTTATGATC |
| KIT c.2447 Forward_2 (Primer C2) | GCCATCATGGAGGATGACGAG |

**Table E3. Thresholds for quality control filtering in Illumina datasets.**

| Dataset | Number of features | UMIs | Percent mt |
|---|---|---|---|
| Ku812_HMC1.2 | 2000 ≤ Features ≤ 7500 | ≤ 45000 | ≤ 10 |
| CML diagnostic | 300 ≤ Features ≤ 6000 | NA | ≤ 8 |
| CML during treatment | 300 ≤ Features ≤ 6000 | NA | ≤ 8 |

NA = not applicable; UMIs = unique molecular identifiers, mt = mitochondrial

**Table E4: Summary of PacBio long-read sequencing and filtering metrics.**

| **Sample** | **Total number of reads** | **Expected reads per cell before filtering[†]** | **Reads retained after filtering for a single Read1 sequence** | **Reads retained after filtering for a single forward primer** |
|---|---|---|---|---|
| Cell lines[‡] (*KIT* c.1679, *KIT* c.2447, *BCR/BCR::ABL1*) | 9,729,819 | 1,425 | 8,982,160 | 4,983,901 |
| Diagnostic CML (*BCR/BCR::ABL1*) | 3,639,857 | 774 | 3,358,100 | 1,957,642 |
| During treatment CML (*BCR/BCR::ABL1*) | 4,916,476 | 882 | 4,703,778 | 2,839,234 |

[†] The calculations were based on the cells that passed the short-read-based quality control filters: 6826 cells for the cell line dataset, 4703 cells for the diagnostic CML dataset, and 5576 cells for the during treatment CML dataset.

[‡] The sequenced sample of the cell lines consisted of three amplification reactions.

**Table E5. Summary of filtered read counts and sequencing saturation by sample and target.**

| Sample | Reads retained after filtering for a single forward primer | Reads retained after cell barcode-UMI deduplication | Sequencing saturation[†] (%) |
|---|---|---|---|
| Cell lines *KIT* c.1679 | 53,743 | 8,464 | 84.25 |
| Cell lines *KIT* c.2447 | 2,975,996 | 496,999 | 83.30 |
| Cell lines *BCR/BCR::ABL1* | 1,954,162 | 585,349 | 70.05 |
| Diagnostic CML *BCR/BCR::ABL1* | 1,957,642 | 300,238 | 84.66 |
| During treatment CML *BCR/BCR::ABL1* | 2,839,234 | 132,686 | 95.33 |

[†] Sequencing saturation was calculated as 1-([reads after cell barcode-UMI deduplication]/[reads before cell barcode-UMI deduplication]).

**Table E6. Sequences used to identify wild-type and mutant alleles in the long-read data**

| Fragment identified | Sequence<br><br>(Note that the reads are in reverse complement) |
|---|---|
| ***BCR* exon 14** – exon 15 | AGGGTGCAGTACAGAT-**TTGAACTCTGCTTAAA** |
| ***BCR* exon 14** – *ABL1* exon 2 | GCCGCTGAAGGGCTT-**TTGAACTCTGCTTAAAT** |
| *KIT* c.2447 WT | TCTCTGGCTAGACCAAAATCAC |
| *KIT* c.2447 MT | ACTCTGGCTAGACCAAAATCAC |
| *KIT* c.1679 WT | ACAACCTTCCACTGTACTT |
| *KIT* c.1679 MT | CCAACCTTCCACTGTACTT |

MT: mutant; WT: wild-type. Bold font corresponds to *BCR* exon 14, and the red bases correspond to the point mutations in *KIT*.

**Table E7. Genotype annotation based on the long-read sequencing data**

| Genotype in sample | Thresholds on the number of long reads detected |
|---|---|
| c.1679 MT in cell line dataset | MT reads ≥ 2 |
| c.1679 WT in cell line dataset | MT reads = 0 and WT reads ≥ 2 |
| c.2447 MT in cell line dataset | MT reads ≥ 4 |
| c.2447 WT in cell line dataset | MT reads = 0 and WT reads ≥ 4 |
| *BCR::ABL1*$^{+}$ in cell line dataset | *BCR::ABL1* reads ≥ 3 |
| Wild-type *BCR* (e14-15) in cell line dataset | *BCR::ABL1* reads = 0 and WT *BCR* reads ≥ 3 |

MT: mutant; WT: wild-type

### Table E8: Comparison of the present method with LOTR-seq

| | Present method | LOTR-seq |
|---|---|---|
| **Target enrichment method** | Semi-nested PCR | Hybridization capture followed by universal PCR |
| **Primer/Probe design** | PCR primers upstream of the loci of interest | Gene-specific hybridization probes |
| **Long-read sequencing platform** | PacBio | Oxford Nanopore |
| **Analysis scope** | Focused interrogation of a limited number of targets | Parallel analysis of a larger number of targets (30 genes demonstrated) |
| **Assay optimization and quality control** | Separate PCR reactions for individual targets enable assay optimization and quality control before long-read sequencing | Targets are enriched together, limiting assay optimization and quality control of individual targets before long-read sequencing. |
| **Gene fusion detection** | Yes | Not investigated |
| **Handling of ambient RNA inherent to the 10X Genomics workflow** | Ambient RNA-derived signals were assessed, and target-specific mutation thresholds were established to improve genotyping specificity | No assessment of how ambient RNA impacts mutation status inference was reported |

### Supplementary Figure Legends

**Supplementary Figure E1. Generation and validation of the cell line scRNA-seq dataset.** **(A)** Scatter plot showing the raw counts of hashtag 3 (x-axis) and hashtag 4 (y-axis). Both axes are shown on a log10 scale. Black lines indicate the thresholds used to assign each cell to a cell line. Only cells assigned to one of the two cell lines were included in the downstream analysis. **(B)** Bar plot showing the percentage of cells in each cell line with at least 1 Illumina short read for *BCR*, *ABL1,* or *KIT*. **(C)** Agarose gel showing reverse transcription PCR amplification of *BCR::ABL1* (*BCR* exon 14-*ABL1* exon 2) and wild-type *BCR* (exon 14-exon 15 junction) in the two cell lines (gel colors inverted).

**Supplementary Figure E2. Identification of single cells harboring *KIT* point mutations.** **(A)** Agarose gel of the amplified *KIT* transcripts using primers upstream of c.1679. **(B)** Sanger sequencing chromatogram of the amplification reaction targeting c.1679. The grey arrow represents the Sanger sequencing primer. The region between c.1679 and c.2447 contains an A-rich region. **(C)** Sanger sequencing chromatogram of the amplification reaction targeting c.2447. The grey arrow represents the Sanger sequencing primer. An A-rich region is present. **(D)** Histograms showing the length distribution of the filtered long reads. **(E)** Histograms showing the number of c.1679 mutated (left) and wild-type (right) long reads per cell barcode. The dashed lines represent a threshold of at least 2 reads. **(F)** Histograms showing the number of c.2447 mutated (left) and wild-type (right) long reads per cell barcode. The dashed lines represent a threshold of at least 4 reads. The zoomed-in histograms show only the cells with read counts meeting the threshold. **(G)** Box plots showing normalized *KIT* expression in genotyped and non-genotyped cells at the c.1679 (left) and c.2447 (right) loci. Genotyped cells include both wild-type and mutant cells, whereas non-genotyped cells refer to cells for which a genotype could not be assigned. All cells, regardless of cell line, were included. Differences between groups were assessed using the Wilcoxon rank-sum (Mann-Whitney U) test. Cliff's delta showed that the *KIT* expression was higher in genotyped cells than in non-genotyped cells at both loci. **(H-I)** Bar plots showing the percentage of mutant reads at the indicated positions. For each substitution, the percentage of mutant reads was calculated as MT / (MT + WT), where WT corresponds to the reference nucleotide indicated at each position. The plots are shown on the same scale as Figures 2K-L for comparison. MT, mutant; WT, wild-type. *** p-value < 0.001.

**Supplementary Figure E3. Detection of *BCR::ABL1* and wild-type *BCR* at the single-cell level. (A)** Agarose gel showing the size of amplified molecules after *BCR::ABL1*-targeted amplification. **(B)** Sanger sequencing chromatogram showing the A-rich region detected downstream the forward primer resulted in truncated molecules. The grey arrow represents the Sanger sequencing primer. **(C)** Sanger sequencing chromatogram showing the sequence of Read 1, the bases that correspond to the cell barcode and UMI as well as the polyT region from the amplification reaction. **(D)** Histograms showing the length distribution of the filtered long reads. **(E-F)** Histograms showing the number of *BCR::ABL1* reads (E) and the number of wild-type *BCR* (F) per cell barcode (CB). The black dashed lines represent a threshold of at least 3 reads. The zoomed-in histograms show only the cells with read counts meeting the threshold. **(G)** Box plot showing normalized expression of *BCR* in genotyped and non-genotyped cells. Genotyped cells include the ones that were either wild-type (WT) *BCR* positive or *BCR::ABL1* positive. Cells from both cell lines were included in the analysis. Differences between groups were assessed using the Wilcoxon rank-sum (Mann-Whitney U) test. Cliff's delta showed that the *BCR* expression was higher in genotyped cells than in non-genotyped cells. **(H)** Box plot showing normalized expression of *ABL1* in the genotyped and non-genotyped cells. Cells were classified as genotyped if they were *BCR::ABL1*-positive. Cells from both cell lines were included in the analysis. Differences between groups were assessed using the Wilcoxon rank-sum (Mann-Whitney U) test. Cliff's delta showed that the *ABL1* expression was higher in genotyped cells than in non-genotyped cells. ** p-value < 0.01, *** p-value < 0.001

**Supplementary Figure E4. Technical validation of single-cell detection of *BCR::ABL1* and wild-type *BCR* in chronic myeloid leukemia samples. (A)** Bar plot showing the percentage of cells with at least 1 Illumina short read aligned to either *BCR* or *ABL1*. **(B)** Sanger sequencing chromatograms from the diagnostic (top) and during treatment (bottom) samples that capture Read 1, cell barcodes, UMIs, and the polyT region from the amplification reactions. **(C)** Read length distribution of the filtered long-reads assigned to wild-type *BCR*. **(D)** Read length distribution of the filtered long reads assigned to *BCR::ABL1*. For the sample obtained during treatment, no reads were detected. **(E)** UMAP visualization of the leukocytes from the diagnostic sample highlighting wild-type *BCR* and *BCR::ABL1* double-positive cells identified at different long-read thresholds. **(F-G)** Box plots showing normalized expression of *BCR* and *ABL1* in genotyped and non-genotyped cells from the diagnostic sample. For the *BCR* plot (F), cells were classified as genotyped if wild-type (WT) *BCR* or *BCR::ABL1* was detected with at least 1 read. For the *ABL1* plot (G), cells were classified as genotyped if *BCR::ABL1* was detected with at least 1 read. Differences between groups were assessed using the Wilcoxon rank-sum (Mann-Whitney U) test. Cliff's delta showed that the *BCR* and *ABL1* expression was higher in genotyped cells than in non-genotyped cells. **(H)** Box plot showing normalized expression of *BCR* in genotyped and non-genotyped cells from the during treatment sample. Cells were classified as genotyped if wild-type *BCR* was detected with at least 1 read. Differences between groups were assessed using the Wilcoxon rank-sum (Mann-Whitney U) test. Cliff's delta showed that the *BCR* expression was higher in genotyped cells than in non-genotyped cells. No *BCR::ABL1*-positive cells were detected in the during treatment sample. ** p-value < 0.01, *** p-value < 0.001
